# Lyophilization induces alterations in cryptococcal exopolysaccharide resulting in reduced antibody binding

**DOI:** 10.1101/2022.03.04.483041

**Authors:** Maggie P. Wear, Audra A. Hargett, John E. Kelly, Scott A. McConnell, Conor J. Crawford, Darón I. Freedberg, Ruth E. Stark, Arturo Casadevall

## Abstract

The structural, antigenic, and immunological characterization of microbial polysaccharides requires purification that often involves detergent precipitation and lyophilization. Here we examine physicochemical changes induced by lyophilization on exopolysaccharide (EPS) of the pathogenic fungus *Cryptococcus neoformans*. Solution ^1^H NMR reveals significant anomeric signal attenuation following lyophilization of native EPS while ^1^H ssNMR shows few changes, suggesting diminished molecular motion and consequent broadening of ^1^H NMR polysaccharide resonances. ^13^C ssNMR, dynamic light scattering, and transmission electron microscopy show that, while native EPS has rigid molecular characteristics and contains small, loosely packed polysaccharide assemblies, lyophilized and resuspended EPS is disordered and contains larger dense rosette-like aggregates, suggesting that structural water molecules in the interior of the polysaccharide assemblies are removed during extensive lyophilization. Importantly, mAbs to *C. neoformans* polysaccharide binds the native EPS more strongly than lyophilized EPS. Together, these observations argue for caution when interpreting the biological and immunological attributes of polysaccharides that have been lyophilized to dryness.

## Introduction

*Cryptococcus neoformans* is protected from the environment and in mammalian infection by a complex polysaccharide capsule. This capsule is a highly hydrated structure and as such, it has a refractive index that is very similar to water, making it difficult to visualize. In the environment, the capsule protects the fungal cell from amoeba predation and dehydration (1, 2). During mammalian infection, the capsule protects the fungal cell from phagocytic cells (3). Additionally, during cryptococcal infection large quantities of cryptococcal polysaccharide are shed into tissue, and this material interferes with effective immune responses (4, 5), overall exacerbating the infection. Detection of cryptococcal polysaccharide in blood and cerebrospinal fluid also provides physicians with important diagnostic and prognostic information for *C. neoformans* disease.

The last five decades have witnessed significant efforts to understand the cryptococcal capsular architecture and yielded important biophysical, chemical, and structural information about the polysaccharide capsule. The dominant polysaccharide component of the *C. neoformans* capsule is glucuronoxylomannan (GXM). Cryptococcal EPS structure has been inferred from light scattering analysis of shed exopolysaccharide (EPS), revealing GXM to be large dense branched polymers (6) that self-aggregate (7) to form dendrimer-like structures (6–8) 1,700-7,000 megadaltons in size (8). The GXM polymer consists of an α-(1,3)-mannose backbone with a β-(1,2)-glucuronic acid branch at every third mannose and varied β-(1,2)- and β-(1,4)-xylose branches from the mannose backbone (9–11). The varied xylosylation results in trimannose repeat motifs, seven of which have been described for GXM (12). In previous studies, the cryptococcal EPS has been isolated using purification steps that require precipitation with cetyl trimethylammonium bromide (CTAB) detergent followed by ethanol precipitation, ultrasonication, dialysis, lyophilization, and base treatment to remove O-acetylation (12). The arrangement of these trimannose motifs into higher order polysaccharide structures has largely remained beyond the reach of technologies for polymer purification and analysis. Changes to the overall polysaccharide organization and structure, depending upon the preparation technique, were evidenced by Circular Dichroism (CD) peaks of higher molar ellipticity in the far-UV region (13). Additionally, ultrafiltration without lyophilization resulted in 14-fold less dense preparations than CTAB precipitation and lyophilization, suggesting that ultrafiltered EPS is organized differently from CTAB-precipitated and lyophilized samples (7). A recent publication postulated that all natural polysaccharides may have physicochemical differences depending upon the method of preparation and that these physicochemical differences translate into functional effects (14).

Here we present evidence of physicochemical alterations to cryptococcal EPS induced by lyophilization to the point of dryness, a technique relied upon for non-sterile EPS isolation. Solution ^1^H NMR spectra of native EPS contain peaks in the SRG region (5.0 – 5.4 ppm) are consistent with GXM; whereas after the sample was lyophilized to dryness and solvated with water, peaks in the SRG region were significantly attenuated or lost. However, magic-angle spinning solid-state ^1^H NMR (MAS ssNMR) spectra indicate that the native EPS and lyophilized samples contained similar material, suggesting that the attenuation of solution NMR signals originates from a change in physicochemical properties rather than changes to chemical structure. In support of this hypothesis, contrasting physical measurements showed that the lyophilized EPS differed from the parent native material in several ways: lyophilized EPS is larger, more mobile, more disordered, and was less reactive with mAbs to GXM. Together, our findings implicate alterations after lyophilization of these polymers. As the majority of published studies on *C. neoformans* EPS rely on lyophilized material, it is essential to consider the impact of these observed differences on the interpretation of previous structural and immunological studies and the design of future investigations that can deepen our understanding of the role of these PS structures in fungal infection.

## Results

### NMR signals are attenuated or absent in rehydrated exopolysaccharide samples

Samples of *C. neoformans* (H99) whole exopolysaccharide (wEPS) were processed only by sterile filtration (0.22μm). We refer to this sample as *native*. Half of this native sample was then lyophilized (~5 d) until the dry weight did not change and solvated with water. We weighed the sample before (97.80 g, average n=3) and after (0.82 g, average n=3) lyophilization. The loss of mass as a result of lyophilization is (average n=3, 96.79 g) 99.16% of the total mass. This result is consistent with a previous study using γ-irradiation to strip the outer capsule, which reduced the cell pellet volume by 85%, suggesting the majority of capsular polysaccharide mass is water (15). Following this analysis, both samples were examined by 1D ^1^H NMR in solution. The solution ^1^H NMR spectrum of the native sample showed a peak set in the structural reporter group (SRG) region (5.0-5.4 ppm), as defined by Cherniak and colleagues (12) (Figure 1A). However, when we examined the same material that had been lyophilized and solvated with water, we found that not all material went into solution. Additionally, the peaks in the SRG region were significantly diminished in intensity or were lost (Figure 1B) (12), even after attempting to re-solubilize the EPS at 37 °C for 14 days with agitation (Figure 1C). While not quantitative due to a lack of baseline peak resolution, overlays of the three spectra, normalized by setting the DSS signal to 1.0, demonstrated that peaks in the SRG region at 5.35, 5.22, and 5.18, were reduced by approximately 60, 80, and 30% respectively, in the lyophilized sample (Figure 1D). Interestingly, while the SRG region in spectra taken from two distinct biological isolates of H99 EPS treated the same way contain the same peak set, the peak intensities differ between samples (Figure S1). However, both sample sets show decrease in peak signal intensity after lyophilization. While there seems to be a greater level of diversity in the polysaccharide of H99 than observed for other strains, the reduction in signal was consistent between samples. *C. neoformans* EPS is generally understood to be solvated with water but we wondered if more hydrophobic solvents could reconstitute the lyophilized material more effectively. We attempted to recover the missing NMR resonances by dissolution of lyophilized wEPS in acetonitrile and dimethyl sulfoxide but neither was superior to water at restoring the signals of the SRG region.

**Figure 1:**
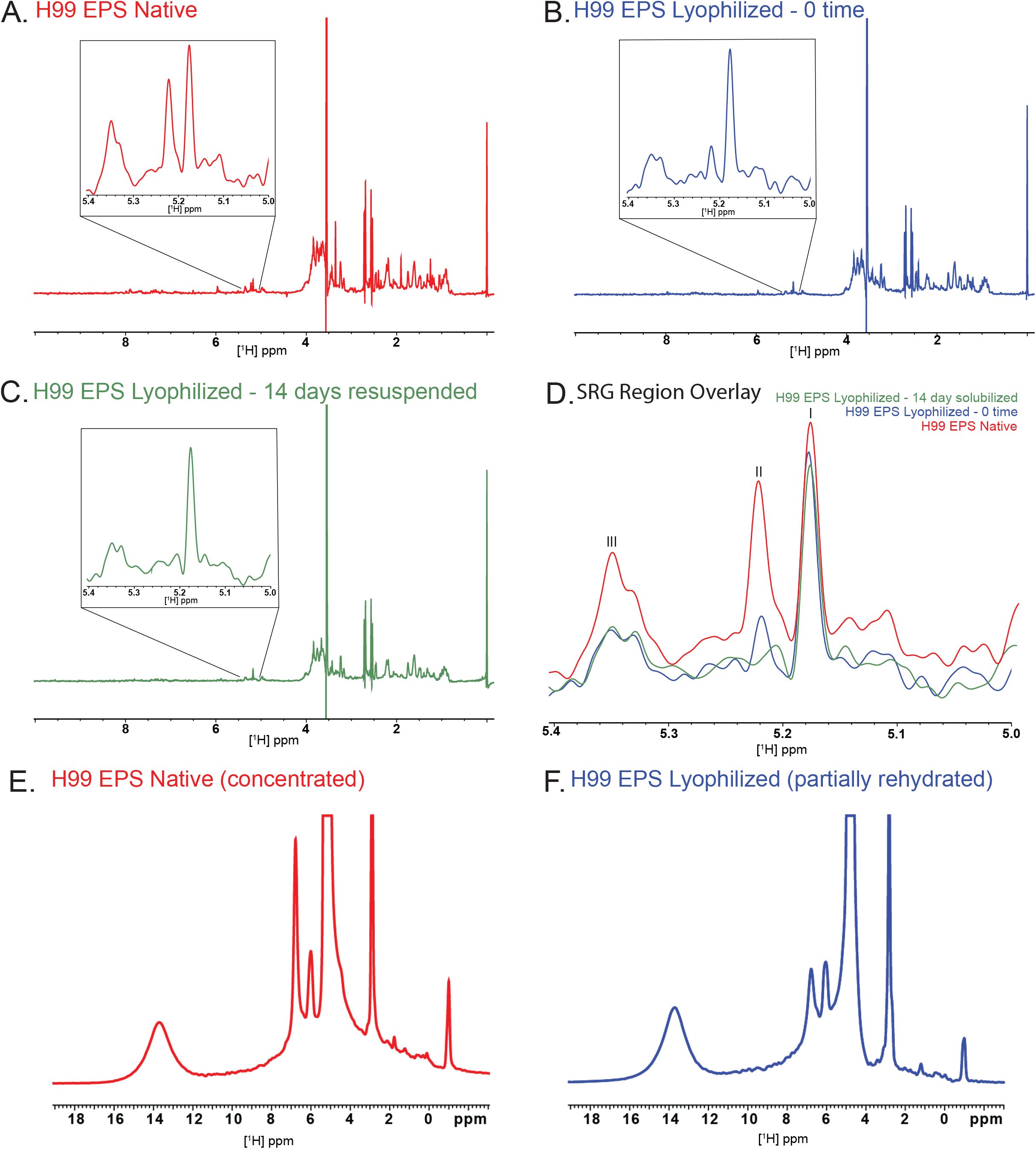
Effects of lyophilization on NMR signals of *C. neoformans* EPS. One-dimensional ^1^H solution NMR spectra and insets expanded vertically by factors of 10 at 60 °C for a native (A) preparation compared with preparations which were lyophilized and solvated with water at time 0 (B) and after 14 days (C); the three spectra are overlayed in (D). SRG region peaks which were integrated indicated as I, II, and III as the motif they belong to is unknown. Peak integrals for the SRG region of the solution-state spectra were compared by setting the respective DSS signals to 1.0. One-dimensional ^1^H solid-state NMR (ssNMR) spectra obtained at room temperature with 15-kHz magic-angle spinning are shown for native (E: concentrated, partially dehydrated) and lyophilized (F: partially rehydrated) samples, normalized according to sample mass. The chemical shifts of the solution- and solid-state spectra were referenced to DSS at 0.0 ppm and water at 4.8 ppm, respectively. The sharp peaks in the ssNMR at 2.9 and 4.8 ppm are attributed to glycine and water, respectively.

### Solid-state ^1^H NMR displayed the same chemical reporter groups in native or lyophilized EPS

We then turned to solid-state ^1^H NMR accompanied by magic-angle spinning (MAS) for partially hydrated wEPS samples, endeavoring to average the orientation-dependent chemical shift tensors to their liquid-state values and remove ^1^H-^1^H dipolar couplings between pairs of nuclear spins that are situated within ~1 nm of one another (16). TThe ssNMR samples were made by (a) concentrating a native EPS solution to a consistency resembling cookie dough and (b) resuspending dry lyophilized EPS with the quantity of water to match that remaining in (a). 1D ^1^H MAS ssNMR showed that both the native (concentrated, partially dehydrated) and lyophilized (partially rehydrated) H99 wEPS samples display the same set of resonances (Figures 1E, F), though the relative peak intensity for the 6.8-ppm signal is notably altered and the SRG signals are overlapped by the solvent in both preparations. These observations suggest that most of the wEPS material present in the resuspended samples maintained its chemical structure but was insufficiently solvated to allow the molecular moieties to become more mobile and thereby more easily observable in the solution-state NMR spectra. In both the native (concentrated) and lyophilized (partially rehydrated) samples, wEPS was solvated sufficiently to be observed by NMR when MAS was used to average out many of the anisotropic spin interactions described above. To our knowledge, these are the first NMR findings that explore the impact of dehydration-rehydration procedures on cryptococcal polysaccharide structure.

### Solid-state ^13^C NMR reveals differences in molecular mobility of polysaccharide in the wEPS samples

A confirmation of the physicochemical rationale for the ^1^H NMR observations and a more detailed comparison of the native (concentrated) and lyophilized (partially rehydrated) EPS materials were available from a follow-up set of ^13^C ssNMR experiments. To probe the impact of hydration at particular molecular sites of the EPS polymers, we acquired both cross polarization magic angle spinning (CPMAS) ^13^C ssNMR (to favor detection of rigid and protonated polymeric moieties) and direct polarization magic angle spinning (DPMAS) ^13^C ssNMR with a short (2-s) delay between successive spectral acquisitions (to ensure inclusion of mobile and disordered chemical groupings in the spectra). Whereas the CPMAS spectra display no EPS signals for either partially dehydrated or partially rehydrated samples (Figure S2), the DPMAS spectra (Figure 2) reveal relatively sharp resonances from the mobile glycan groups (~62-105 ppm) in both native (concentrated) and lyophilized (partially rehydrated) samples, but no significant contributions from alkene or carboxyl carbons with chemical shifts above 110 ppm.

**Figure 2.**
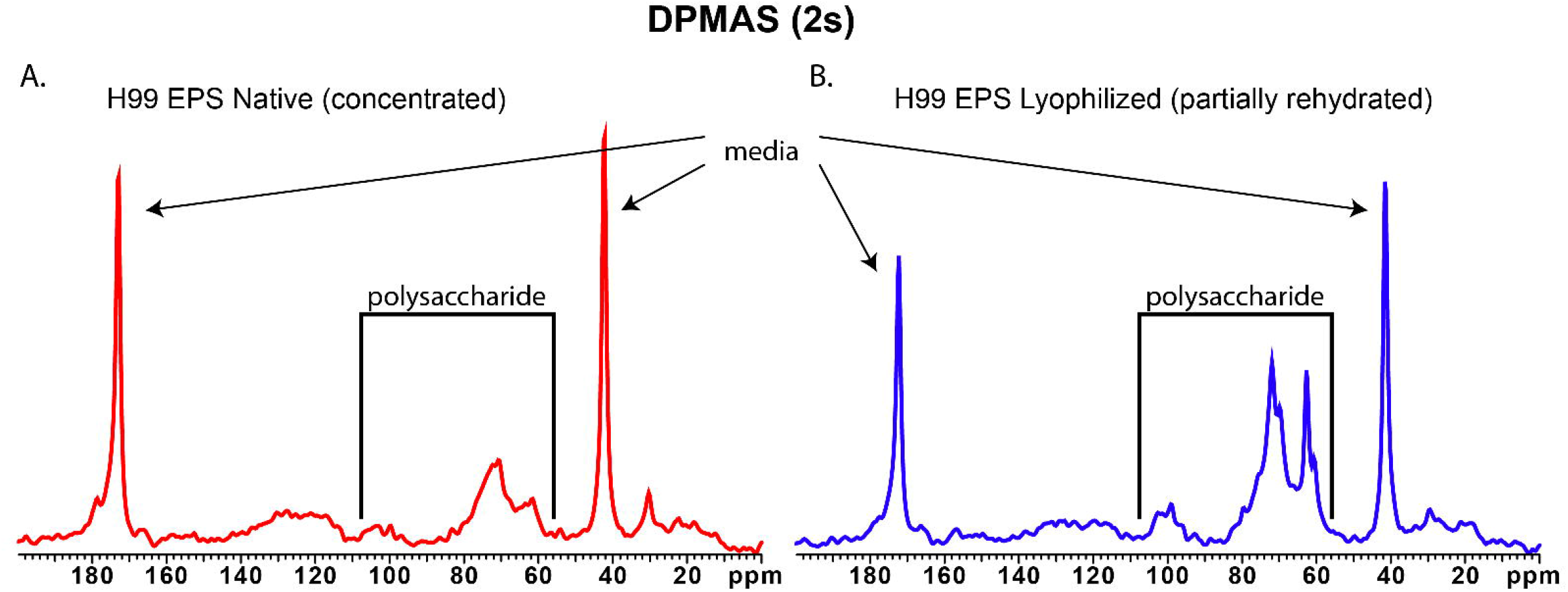
Effects of lyophilization on solid-state ^13^C NMR spectra of EPS. 150 MHz ^13^C NMR spectra of *C. neoformans* EPS samples obtained with 15 kHz magic-angle spinning (MAS), comparing samples that were native (partially dehydrated) (left) and lyophilized (partially rehydrated) (right). DPMAS experiments with short (2 s) delays between successive cycles of signal acquisition, favoring carbon moieties that tumble rapidly in many directions. Sharp resonances at 40 and 170 ppm are attributed to glycine in the culture media. (No EPS signals are observed in CPMAS experiments that favor rigid carbon moieties with nearby hydrogens.)

Notably, the major glycan resonances between ~62 and 105 ppm are sharper and thus better resolved in the lyophilized (partially rehydrated) sample, indicating more complete solvation and motional averaging of the polysaccharide structures. The mobility that yields resolved ^1^H and ^13^C NMR spectra under magic-angle spinning acquisition conditions can be attributed to the hydrophilic nature of the sugar ring structures.

### Electron Microscopy of EPS shows rosette-like assemblies in rehydrated lyophilized sample

To further investigate the effects that lyophilization might have on EPS, we turned to Transmission Electron Microscopy (TEM) to examine the architecture of the EPS samples. This analysis shows that the native EPS is less dense and contains vesicles (Figure 3A). The presence of vesicles is not surprising since these are shed by *C. neoformans* during capsule growth (17, 18) and would be retained by the filtration step. In contrast, no vesicles were observed in the lyophilized and reconstituted samples, possibly reflecting collapse of these structures during the drying procedure (19). The lyophilized and reconstituted material does contain dense, rosette-like assemblies, similar to those observed previously for cryptococcal capsular polysaccharide isolations and glycogen (Figure 3B) (6, 20).

**Figure 3:**
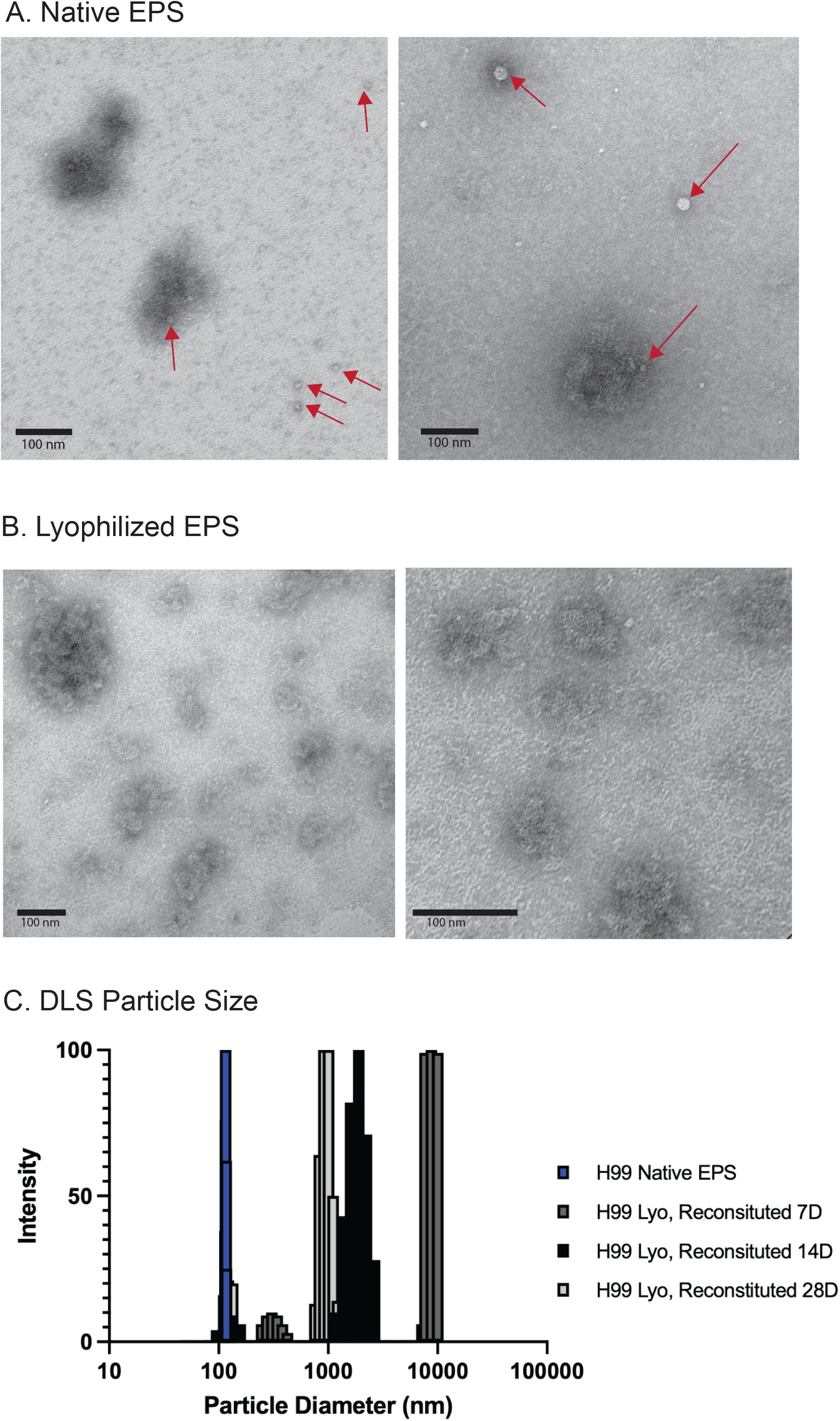
*C. neoformans* EPS undergoes biophysical changes over time in solution. Native EPS (A) and lyophilized and reconstituted EPS (B) samples were examined by transmission electron microscopy (TEM) with negative staining. Native EPS contains a few aggregates as well as extracellular vesicles (EVs), while lyophilized and resuspended EPS contains many dense rosette-like structures previously reported for *C. neoformans* polysaccharides and glycogen (6). Native EPS and lyophilized and reconstituted samples kept in solution for 7, 14, or 28 days were examined by DLS (C). Lyophilized and resuspended samples have a larger particle size than native samples, judged by autocorrelation intensity of the scattered light as a function of particle diameter.

### Dynamic light scattering shows size differences as a function of solubilization time

Dynamic light scattering (DLS) revealed that the average effective diameter for particles in the native wEPS preparation were ~115 nm but after lyophilization these increased in size (~300 nm and ~8500 nm) and cover a wider size range (Figure 3C), consistent with reported sizes for EPS particles (6–8). Over the course of 28 days in solution (D_2_O), the effective diameter decreased (~950 nm with smaller particles) (Figure 3C).

### ELISA uncovers antigenic differences between native and lyophilized EPS

To examine how native and lyophilized EPS are bound by monoclonal antibodies (mAbs) to GXM, we performed capture ELISA, a standard assay for determining GXM concentration in a sample (21). Figures 4A and 4B show that mAbs to GXM binds more strongly to native than lyophilized wEPS. This finding is consistent with previous observations comparing CTAB- to ultrafiltration-prepared EPS, wherein ultrafiltered EPS samples bound both 12A1 and 18B7 better than CTAB precipitated EPS in direct ELISAs (7). These results suggest that there are more available epitopes in the native polysaccharide than in wEPS that has been lyophilized and resuspended.

**Figure 4:**
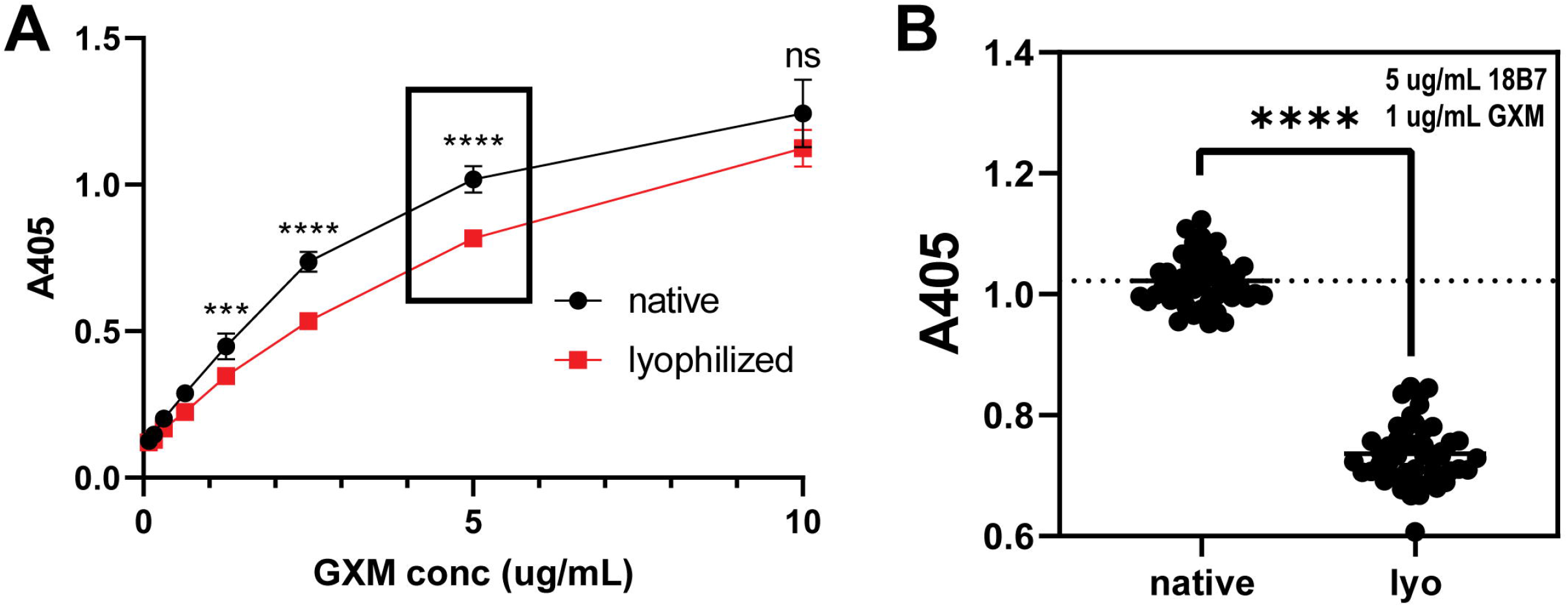
Lyophilization of *C. neoformans* EPS alters biological functions. Native EPS and lyophilized and resuspended samples were assayed for anti-GXM mAb binding by capture ELISA. Binding curves (A) of serially diluted mAb 18B7 as a function of EPS concentration in capture ELISA. Native EPS generally binds more strongly to the 2D10/18B7 capture/detection mAb pair than lyophilized EPS at a given mAb/antigen concentration. Statistical analysis (B) of binding in a capture ELISA double-array assay varying both EPS and mAb concentration shows that native EPS is statistically significantly better bound by anti-GXM mAbs than lyophilized EPS. **** p-value < 0.0001.

### Proposed model of dehydration-rehydration effects

Previous work by Cordero *et al*. showed that both EPS and CPS that were lyophilized and rehydrated had hydrodynamic properties consistent with a dendrimer-like conformation (6). Similar aggregates can be visualized on the surface of cryptococcal cells after dehydration imaged by scanning electron microscopy (SEM) and as secreted particles by transmission electron microscopy (TEM) (6). These dendrimer-like formations have a higher density at the core and more dispersed radial polymers. CTAB-precipitated EPS is also more dense (14-fold) than the ultrafiltered native material (7). When these observations are considered in light of the observations presented here, the suggested dendrimer-like formations for GXM are consistent with our lyophilized and resuspended material, but not with native materials (Figure 5), though other forms cannot be ruled out. Prior to lyophilization, GXM polymers shed into solution as EPS by *C. neoformans* are small, rigid, ordered, and hydrated. Lyophilization results in the adoption of mobile, disordered, dense, and aggregated architectures. While more time in solution may eventually restore these polymers to resemble the native polysaccharide more closely, the monthlong incubation time used in our study was insufficient to return them to their native state (Figure 5).

**Figure 5:**
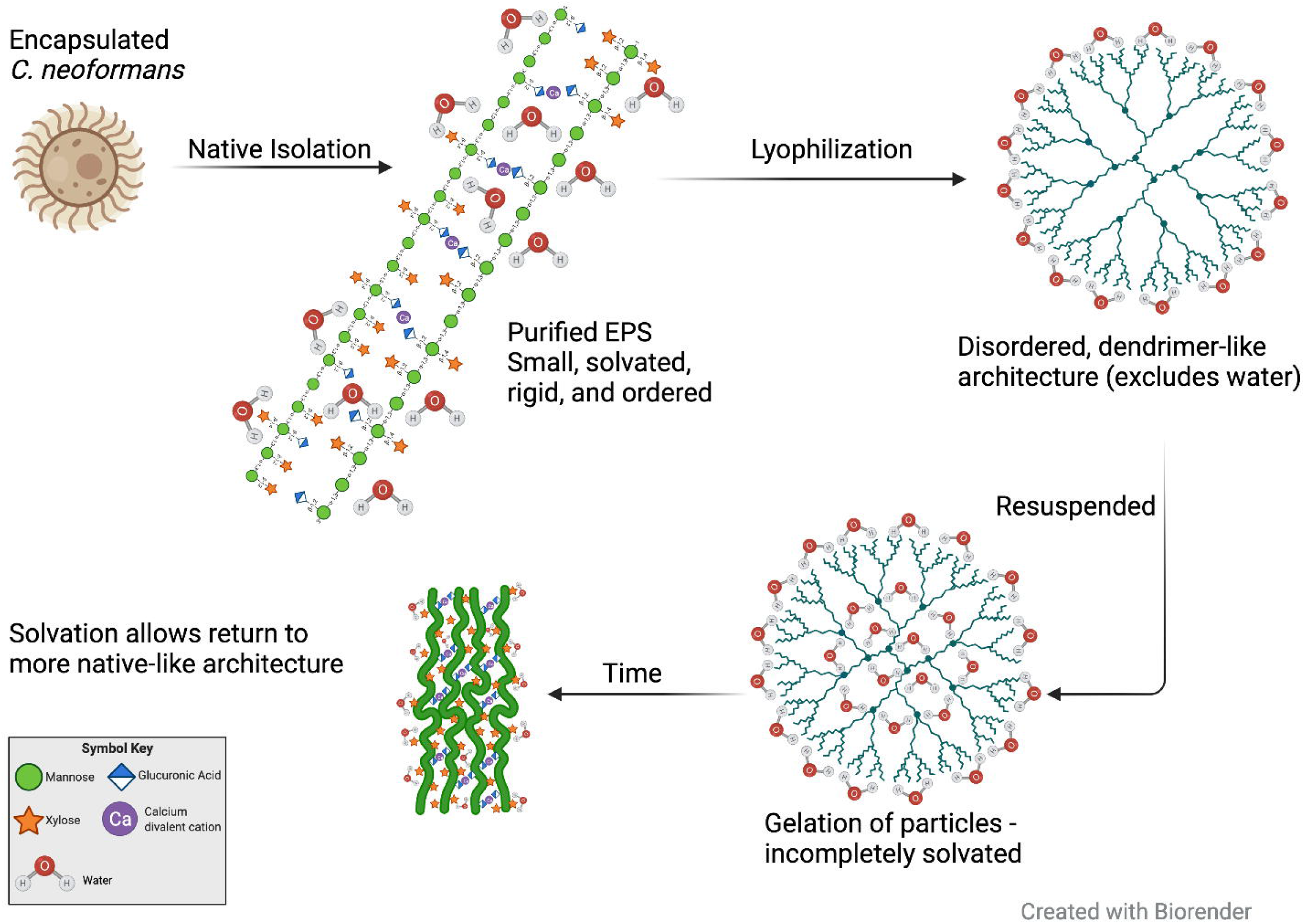
Structural model for the effects of lyophilization on *C. neoformans* EPS structure. EPS harvested from encapsulated *C. neoformans* is small, hydrated, rigid, and ordered when purified in its native form. Lyophilization causes alterations to the native form, resulting in disordered, dendrimer-like conformations that exclude water. Solvation of the lyophilized sample proceeds through gelation, wherein particles aquire a hydration shell but not structural waters. Over time in solution (28+ days) the dendrimer-like conformation will be lost and a polymer-like structure more similar to, but not the same as, the native structure is adopted.

## Discussion

Historically, analysis of the cryptococcal capsule has relied on the examination of the shed EPS polymers. These analyses indicate that EPS is composed of large, dense, branched polymers (6–8). Techniques for the isolation of EPS have evolved since these initial analyses, however, the maintenance of sterile sample preparations is challenging, resulting in the use of lyophilization for long term polysaccharide sample storage. There are indications in the literature that cryptococcal polysaccharide is altered by the method of isolation (7, 13, 22). Light scattering measurements of EPS samples reveal nine-fold larger particles when precipitation with the cationic detergent CTAB is utilized compared to isolation by ultrafiltration show (7), (13). Some investigators have stated outright that the structure of EPS varies by method of preparation as well as that CTAB isolation alters the secondary structure of polysaccharide (22). It is of utmost importance to define the impact of polysaccharide preparation on the physicochemical and antigenic properties of EPS, as our understanding of the immunoregulatory role of cryptococcal EPS is largely derived from analysis of purified polysaccharide on immune cells.

One of the functions of the capsule is to protect the fungal cells from dehydration (1). GXM, the predominant polysaccharide of the cryptococcal capsule, derives its hydrophilic nature from its components – mannose, xylose, and glucuronic acid – as well as the water coordination necessary to maintain the divalent cation bridges formed between glucuronic acid residues (7). To appreciate the necessity of water to both the capsule and its composite polymers, we can examine the sheer quantity of water present. As noted above, when samples are weighed prior to and after lyophilization, water makes up a significant proportion of the mass (98% in this study and 85% in the gamma irradiation study). The additional data presented here, including EM, DLS, solution NMR and solid-state NMR imply that there may be internal or structural water molecules that are necessary for the overall structure and organization of the polysaccharide assembly. This suggests that water is critically important to the three-dimensional structure of cryptococcal polysaccharide, not only forming a hydration shell, but including structural waters. Our observations indicate that after lyophilization the polysaccharide can be partially solvated (hydration shell) but does not allow for the incorporation of these structural waters.

In this work, we examine an observed difference between native and lyophilized EPS samples in which we noted the attenuation of anomeric carbon signals in ^1^H solution-state NMR after lyophilization. However, ssNMR ^1^H spectra show no significant difference between the signals in the native and lyophilized samples, suggesting that the signal attenuation in solution was due not to a chemical change, but to incomplete solvation and subsequent failure to restore the polysaccharide assembly to its native state. The physiochemical alterations resulting from loss of water could include increased molecular size, decreased molecular mobility, and limited angular excursions, which in turn would enhance nuclear spin relaxation and broaden the resonances to the point that individual signals would appear to vanish. Further comparison of the two samples by TEM and DLS indicates that lyophilized and resuspended samples are larger, more mobile, and disordered. While most of these observations are expected, the increased mobility runs counter to the dendrimer-like aggregation observed in the lyophilized sample. We would note that the ^13^C ssNMR of lyophilized (partially rehydrated) samples exhibit greater flexibility than the native (concentrated) samples, but neither of these states exhibits the rapid isotropic motions that would yield well-resolved solution-state NMR spectra.

Previous reports suggest that inter-polymer interactions occur through divalent cation bridges formed between glucuronic acid residues of independent polymer. One interpretation of this effect is that the solvation of these divalent cation bridges proceeds slowly over time. However, we did not observe lyophilized molecules returning to their pre-lyophilization, native, state after a month in solution. This may be due to incomplete hydration, loss of mannose O-acetylation, or incomplete solvation wherein some polymers are recalcitrant to reconstitution. It is possible that the application of other conditions such as higher temperature and/or different solution conditions (pH or electrolyte concentrations) could return the polymers to their native state.

Although these physicochemical alterations to cryptococcal EPS are interesting in their own right, we also observed functional changes as suggested by Yi *et al* (14). *C. neoformans* EPS has been shown to mediate numerous deleterious effects on host immune function (23), which presumably result from the interaction of carbohydrates with cellular receptors. Alterations in antigenic properties can be inferred from differences in mAb reactivity observed by capture ELISA. Our observed reduction in binding of the lyophilized sample by capture ELISA suggests decreased epitope prevalence and/or accessibility, revealing that the physicochemical alterations effected on the polysaccharide by lyophilization have a functional impact. Antibody interactions may require a specific polysaccharide arrangement that is altered by lyophilization, suggesting the need to revisit these observations with native material. Interestingly, although mAbs 2D10 and 18B7, the capture and detection mAbs used in this experiment, respectively, were raised against CTAB-prepared GXM conjugates (which are lyophilized), they preferentially bind to native wEPS. This trend may reflect enhanced antigen presentation in the smaller native EPS particles. Furthermore, it is possible that similar effects occur when other microbial polysaccharides are isolated by precipitation, lyophilization and reconstitution techniques, which argues for caution in extrapolating observations with different methods of preparation to those present in native macromolecules. In a recent review, Yi and colleagues discussed the dehydration of polysaccharides as one of the key procedures in processing them and noted that vacuum drying and hot air drying both lead to larger molecular weights, poorer solubility, and increased incidence of aggregation compared to freeze drying (14). We have observed each of these three effects upon lyophilization of *C. neoformans* EPS. Similarly, increased apparent size by DLS, poor solubility, and increased aggregation have also been reported for polysaccharides from acorn (24), Chinese medicinal herb *Bletilla striata* (25), mushroom *Inonotus obliquus* (26), comfrey root (27), and finger citron fruits (28). Further studies will be necessary to tease apart the effects of isolation, freeze-, and vacuum-drying on polysaccharides, particularly for cryptococcal EPS. Nevertheless, our observations together with reports of other polysaccharides undergoing physicochemical alteration upon dehydration (14) suggest that this may be a widespread phenomenon for such polymers and argues for caution when interpreting findings from rehydrated material.

In conclusion, scientists investigating the immunological properties of cryptococcal polysaccharides should be aware that the method of purification can affect its physicochemical properties, which in turn can affect some of the immunological properties of polysaccharides. The physicochemical alterations exacted by CTAB and lyophilization upon polysaccharides could explain much of the variability in published studies (29–32) and suggest the need for a renewed effort to characterize cryptococcal polysaccharides using isolation techniques that maintain these molecules in their native states.

## Experimental Procedures

### Fungal Growth and Exopolysaccharide Isolation

*C. neoformans* serotype A strain H99 (ATCC 208821) cells were inoculated in Sabouraud rich medium from a frozen stock and grown for two days at 30° C with agitation (150 rpm). Capsule growth was induced by growth in chemically defined media (7.5 mM glucose, 10 mM MgSO4, 29.4 mM KH_2_PO4, 6.5 mM glycine, and 3μM thiamine-HCl, pH 5.5) for 3 days at 30°C, with agitation (150 rpm). The supernatant was isolated from cells by centrifugation (4,000 × g, 15 min, 4°C) and subsequently sterilized by passing through a 0.45 μm filter. Native samples were concentrated while lyophilized samples were freeze dried to complete dryness, defined by no change in mass with time, for an average of 5 to 7 days.

### Solution NMR

1D ^1^H NMR data were collected on either of two spectrometers: a Bruker Avance II (600 MHz), equipped with a triple resonance, TCI cryogenic probe and Z-axis pulsed field gradients or a Bruker Avance III HD (700 MHz), equipped with an XYZ gradient TCI cryoprobe. Spectra were collected at 60°C, with 64 scans and a free induction decay size of 84336 points. Standard Bruker pulse sequences were used to collect the 1D data (p3919gp and zggpw5). Data were processed in Topspin (Bruker version 3.5) by truncating the FID to 8192 points using a squared cosine bell window function and zero filling to 65536 points.

Lyophilized samples were dissolved in deuterated water to a concentration of 50 mg/mL or greater. Native samples were diluted by adding 300 μl of D_2_O to 200 μl of sample. All NMR samples contained DSS-d_6_ for chemical shift calibration and peak intensity comparisons.

### Dynamic Light Scattering

Measurement of EPS particles by DLS was performed with a Zeta Potential Analyzer instrument (Brookhaven Instruments). The particle sizes in the suspension were measured for native samples as well as lyophilized and rehydrated samples at different time points during a period of 28 days. Data are expressed as the average of 10 runs of 1-min data collection each. The multimodal size distributions of the particles were obtained by a non-negatively constrained least squares algorithm based on the intensity of light scattered by each particle. The multimodal size distributions of particles from each sample were graphed for comparison.

### ELISA

For capture ELISA, Microtiter polystyrene plates were coated with goat anti-mouse IgM at 1 *μ*g/ml (SouthernBiotech, Birmingham, AL) and then blocked with 1% BSA blocking solution. 2D10, a murine anti-GXM IgM, was subsequently added at 10 *μ*g/mL as the capture antibody. Next, lyophilized or native wEPS samples were added to each half of the plate and serially diluted. 18B7, a murine anti-GXM IgG1, was added at 10 *μ*g/mL and serially diluted in the opposite direction as the GXM dilution. The direct ELISA was performed by coating plates directly with antigen (native or lyophilized EPS) at 1 *μ*g/mL, followed by 18B7 at 5 *μ*g/mL to each well. For both, the assays were developed by sequential addition of goat anti-mouse IgG1 conjugated to alkaline phosphatase at 1 *μ*g/mL and 1 mg/mL *p*-nitrophenol phosphate substrate. The absorbance of each well was measured at 405 nm after a short incubation at 37°C. Between each step of the ELISAs, the plate was incubated for 1 h at 37°C and washed three times in 0.1% Tween 20 in Tris-buffered saline.

### Solid-State NMR

Partially dehydrated samples were prepared by lyophilizing a 5-mL EPS solution for 18 hours to obtain 309 mg of a ‘cookie dough’ material that was packed into a 3.2-mm OD ssNMR rotor. Partially rehydrated samples were prepared by adding 0.08 mL of water to 211 mg of fully dried EPS powder, matching the weight percent of the ‘cookie dough’ and yielding 297 mg of a ‘sticky batter.’ NMR spectra were acquired with a Varian (Agilent) DirectDrive2 spectrometer operating at a ^1^H frequency of 600 MHz and using a 3.2-mm T3 HXY Magic Angle Spinning (MAS) probe (Agilent Technologies, Santa Clara, CA). These data were acquired on 34.1 and 39.7 mg, respectively, of concentrated (partially dehydrated) and lyophilized (partially rehydrated) samples using a spinning rate of 15.00 ± 0.02 kHz and a nominal temperature of 25°C. The ^1^H spectra were obtained with a single 90°pulse, whereas ^13^C spectra used either 1-ms ^1^H-^13^C cross polarization (CP) with a 10% ramp of the ^1^H power and 3 s between data acquisition sequences or direct polarization (DP) with a 2-s recycle delay. ^1^H decoupling with a radiofrequency field of 109 kHz was applied during signal acquisition with the small phase incremental alternation method (33). After apodization of the data with a decaying exponential function to improve the signal-to-noise ratio and Fourier transformation, the spectra were referenced to H_2_O at 4.8 ppm.

### Negative staining with Uranyl Acetate and Transmission Electron Microscopy

Samples (10 uL) were adsorbed to glow discharged (EMS GloQube) ultra-thin (UL) carbon coated 400 mesh copper grids (EMS CF400-Cu-UL), by floatation for 2 min. Grids were quickly blotted then rinsed in 3 drops (1 min each) of TBS. Grids were negatively stained in 2 consecutive drops of 1% uranyl acetate with tylose (UAT), then quickly aspirated to get a thin layer of stain covering the sample. Grids were imaged on a Hitachi 7600 TEM (or Philips CM120) operating at 80 kV with an AMT XR80 CCD (8 megapixel).

## Funding and Additional Information

A.C. and M.P.W. were supported by NIAID R01AI152078; S.A.M. was supported by T32 AI00741726. R.E.S., J.E.K., and A.C. were supported for this work by National Institutes of Health Grant R01-AI052733. C.J.C. was funded by the Irish Research Council postgraduate award (GOIPG/2016/998). The 600 MHz ssNMR facilities used in this work are operated by The City College of New York and the City University of New York Institute for Macromolecular Assemblies. The content is solely the responsibility of the authors and does not necessarily represent the official views of the National Institutes of Health.

## Acknowledgments

The mechanism figure was created with Biorender software. We thank Dr. Christine Chrissian for assistance with the ssNMR data acquisition and analysis. We thank Barbara Smith and the Electron Microscopy core facility at Johns Hopkins School of Medicine for their analysis of the EPS samples as well as conversations about these data.

---

**Supplemental Figure 1. Solution ^1^H NMR variation observed between biological replicates of H99 EPS.** Unprocessed *C. neoformans* H99 EPS samples (Native) from two different biological replicates were examined by 1D ^1^H NMR. The sample in red is portrayed throughout this work because the peak set in the SRG region was easier to define than in replicate 2. The same reduction in signal was observed for both samples after lyophilization.

**Supplemental Figure 2. Effects of lyophilization on solid-state ^13^C NMR spectra of EPS.** 150 MHz ^13^C NMR spectra of *C. neoformans* EPS samples obtained with 15 kHz magic-angle spinning (MAS). CPMAS experiments that favor rigid carbon moieties for which cross polarization from nearby hydrogen nuclei is efficient. A. Native H99 EPS (concentrated, partially dehydrated). B. Lyophilized H99 EPS (partially rehydrated).

