## Supplementary figures and images for "Lyophilization induces alterations in cryptococcal exopolysaccharide resulting in reduced antibody binding"

### Supplemental Figure 1

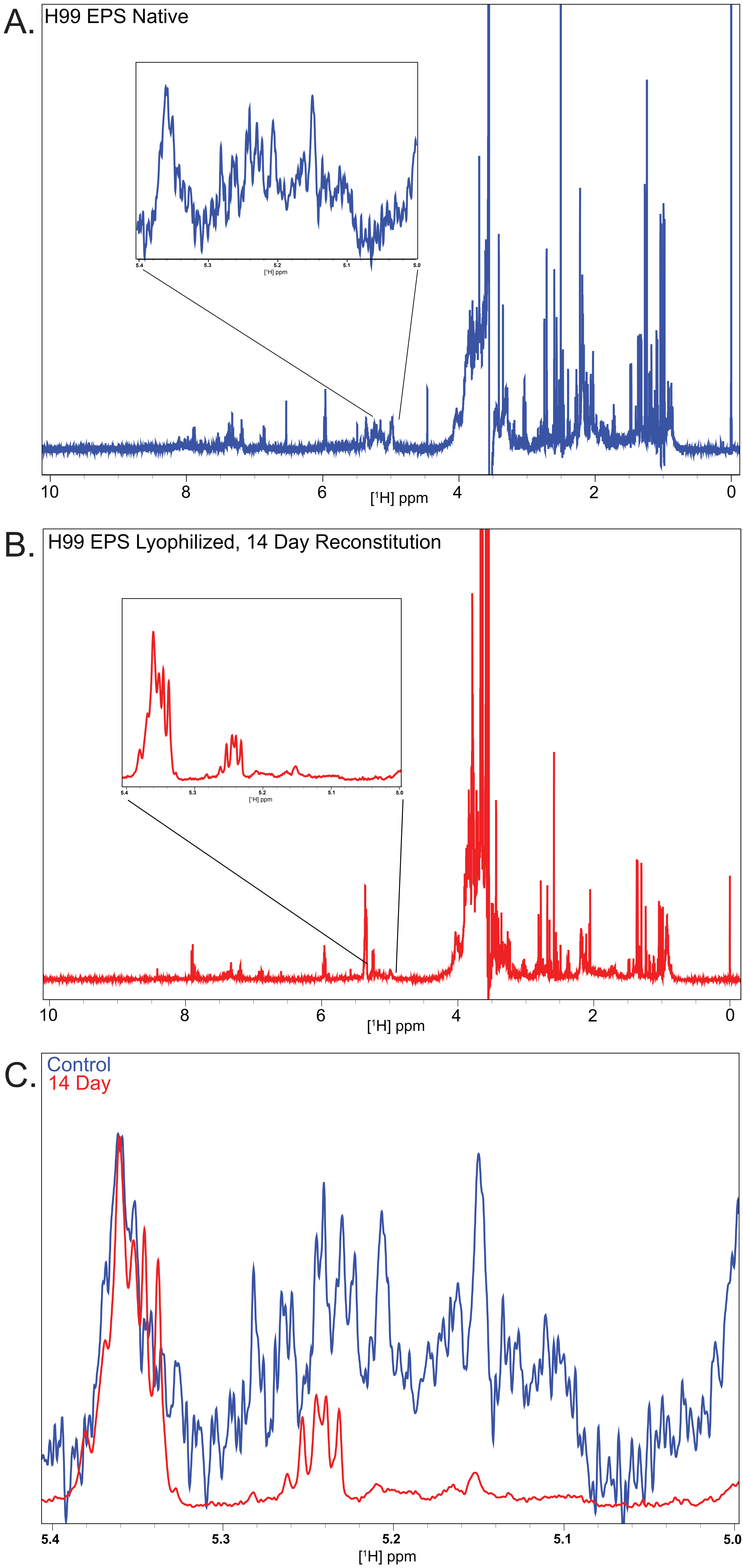

### Supplemental Figure 2

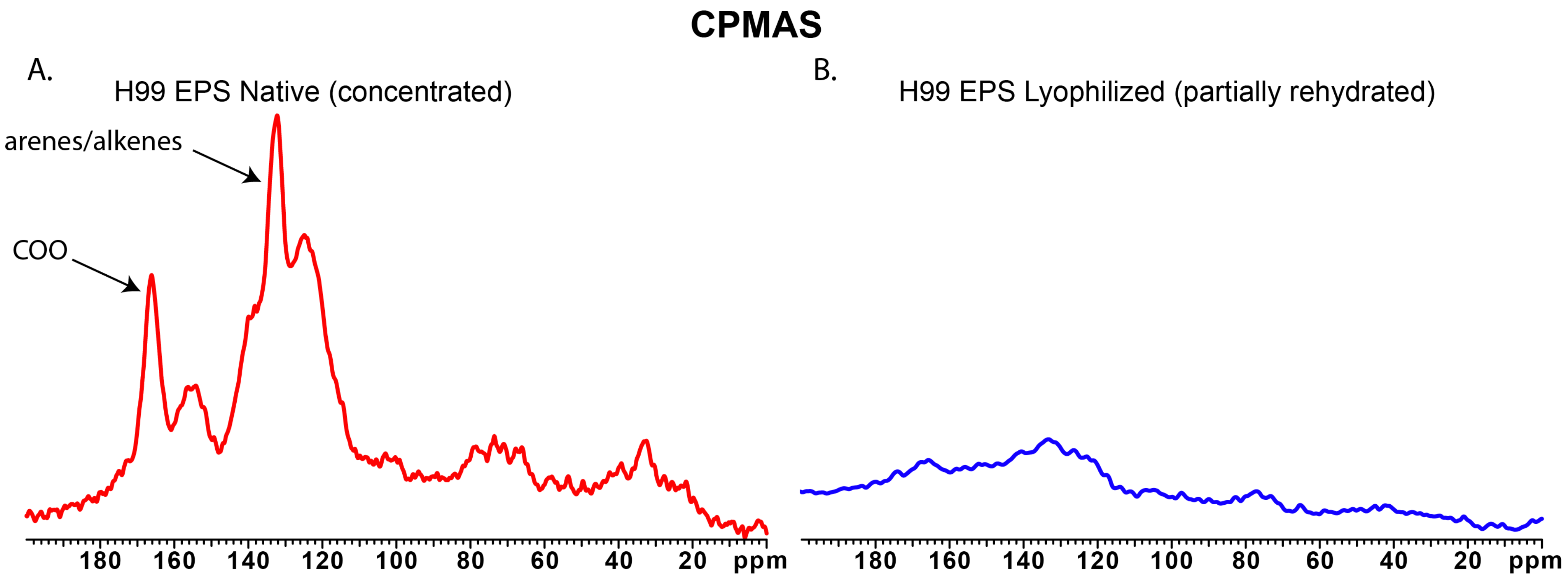
